# Conformational heterogeneity and state-dependent stability of active-site loops in *Trametes versicolor* laccase

**DOI:** 10.1101/2025.11.20.689517

**Authors:** Sudipta Mitra, Ranjit Biswas, Suman Chakrabarty

## Abstract

How can a single active site accept substrates that differ widely in size, shape and charge? We have addressed this question for the fungal laccase from *Trametes versicolor*, a promiscuous multicopper oxidase, by mapping the conformational ensemble of the substrate-free (but fully copper-loaded) enzyme. Combining 50 μs of unbiased atomistic molecular dynamics seeded through tICA-metadynamics with a variational autoencoder and VAMPnets, we resolve a set of metastable states that differ mainly in the conformations of the two loops, L1 (G159–L164) and L2 (F332–F337), lining the substrate binding pocket. The loops interconvert on the sub-millisecond timescale, and their rearrangement is accompanied by an extensive rewiring of the hydrogen-bond network well beyond the active site. Conformations from our earlier dye-bound simulations map onto specific regions of this landscape, showing that the bound poses correspond to conformations that the free enzyme already visits. A correlation network connecting the two loops passes through the protein region surrounding the T1 copper and singles out T114, the residue with the highest betweenness centrality along the path. Nonequilibrium alchemical calculations show that T114A destabilizes the five states by very different amounts, between 3.2 and 8.2 kcal mol^−1^ relative to a common peptide reference, with the largest penalties occurring in states carrying a persistent T114–D118 hydrogen bond. A single side chain interaction, located away from the substrate-binding contacts, thus offers a handle for tuning the conformational balance of the active site, and this prediction can be tested directly by mutagenesis.

**Significance statement:** Enzymes acting on many substrates present a structural puzzle: one active site must accommodate molecules of very different size, shape and charge. Extensive atomistic simulations of a fungal laccase show that the two loops forming the rim of its active site interconvert between several distinct conformations, each with its own hydrogen bonding pattern, and that dye-bound structures match specific members of this pre-existing set. A correlation analysis points to a threonine behind the loops, and free energy calculations show that removing its side chain shifts the relative stability of the conformers by several kcal mol^−1^. The calculations therefore predict that conformational balance at an active site can be tuned from a distance, a prediction directly testable by mutagenesis.

## Introduction

Laccases are multicopper oxidases that couple the one-electron oxidation of a substrate at the mononuclear T1 copper site to the four-electron reduction of molecular oxygen to water at the trinuclear copper cluster (TNC)^1,2^. They are found in fungi, plants, bacteria and insects^3^, and they act on a remarkably wide range of phenolic and non-phenolic compounds^4^. This promiscuity is what makes them attractive for dye decolorization, removal of pharmaceutical contaminants, biosensing and biofuel production^5–8^. It also raises a question that is interesting from a purely physical point of view: how does one binding pocket accommodate molecules that differ so widely in size, shape and charge? Conformational heterogeneity of the enzyme is an obvious candidate^9,10^, alongside the chemical character of the pocket and the redox requirement of the reaction. Our objective in this work is to characterize this heterogeneity in the substrate-free enzyme and to ask whether the balance between the conformers can be perturbed in a controlled and experimentally testable manner.

We have shown earlier that *T. versicolor* laccase interacts with five dye molecules of widely different size, shape and charge^11^: brilliant blue (net charge -2), coumarin 343 (−1), methyl green (+2), crystal violet (+1) and thioflavin T (+1). The accompanying UV–visible measurements follow the loss of the characteristic absorbance of each dye, which is a direct indication of decolorization; identification of the reaction products, and hence the distinction between partial transformation and complete degradation, lies beyond that experiment. Redox mediators are known to extend the range of compounds accessible to laccases, and direct enzyme–substrate oxidation and mediator-assisted oxidation are distinct processes^5^. Previous experimental studies have quantitatively shown that laccase can decolorize brilliant blue, crystal violet and methyl green with or without mediators^12,13^. In the present work we use the dye-bound trajectories of ref. 11 as structural references for comparison with the substrate-free enzyme.

The crystal structure of *T. versicolor* laccase in complex with 2,5-xylidine (PDB 1KYA) provides our starting model^14^. Two loops, L1 (G159–L164) and L2 (F332–F337), form the rim of the substrate binding pocket (Fig. 1)^15^. Four copper ions constitute the catalytic machinery: the mononuclear T1 copper close to the substrate binding site, and the TNC made up of the T2 and the two T3 coppers^2,16^. H458 coordinates the T1 copper and can also interact directly with a bound substrate^17^. Throughout this paper we use the word “apo” in the restricted sense of substrate-free: all four copper ions are present in every simulation reported here, and no cofactor has been removed.

**Fig. 1.**
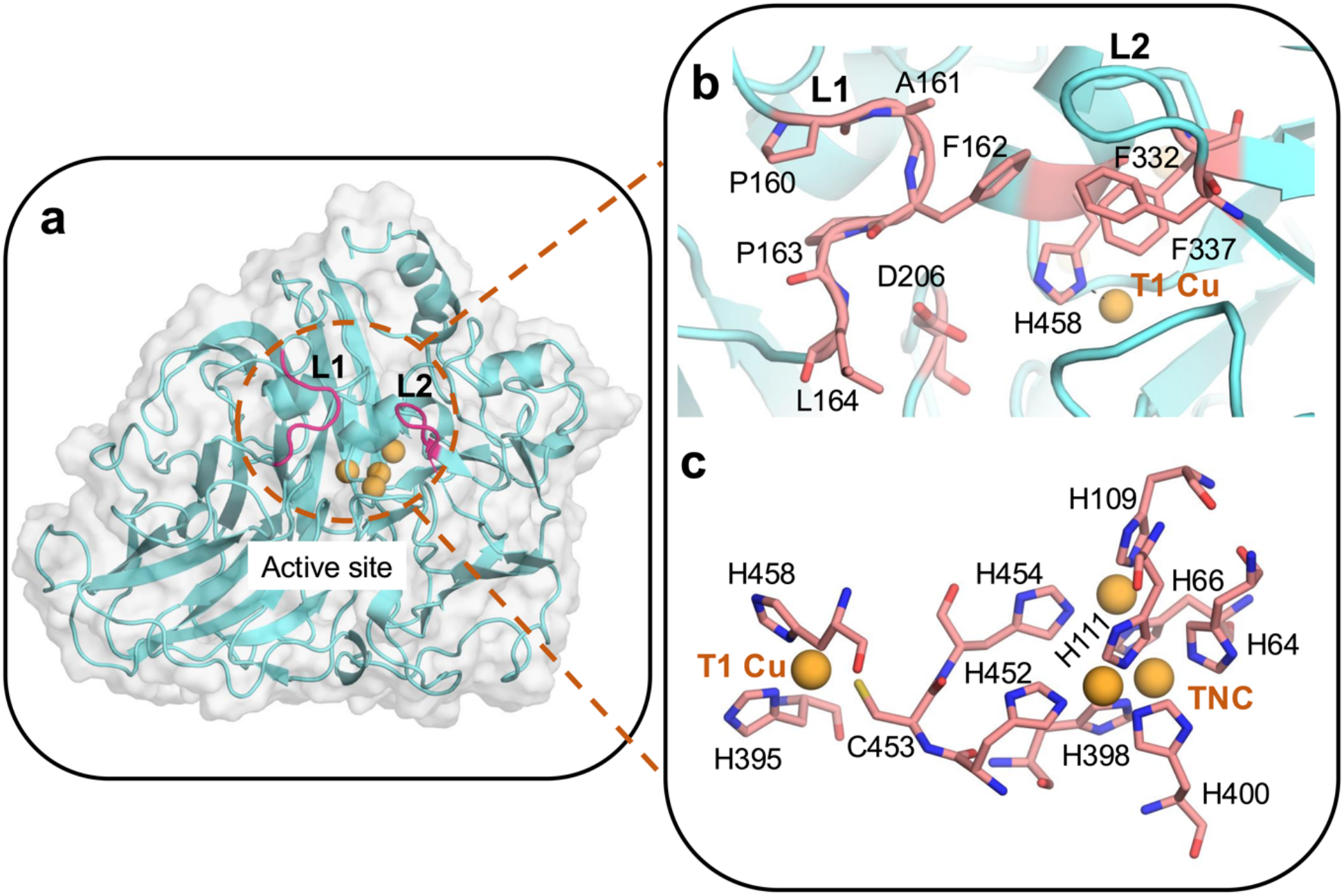
Structure of the *Trametes versicolor* laccase. (a) The overall fold (PDB 1KYA), with the two active site loops L1 (G159–L164) and L2 (F332–F337) highlighted in magenta and the copper ions shown as orange spheres. (b) Close-up of the substrate binding pocket showing the two loops, the pocket residues, and the T1 copper ligand H458. (c) The T1 copper and the trinuclear copper cluster (TNC) with their coordinating residues shown as sticks.

It is well known that mutations both inside and outside the binding pocket can modulate laccase activity^18,19^. In *T. versicolor* laccase, substitutions at F162, F332 and F337 alter the substrate-dependent reactivity^19^, while mutation of D206 shifts the pH profile of the activity^20^, and an additional ABTS binding site has been identified in the bacterial CotA laccase^21^. Observations of this kind motivate our search for interactions that lie outside the immediate substrate-contacting region and are nevertheless coupled to the conformation of the active site.

Substrate-bound crystal structures and simulations of substrate association provide complementary views of laccase recognition^21,22^. Here we take the route of first characterizing the substrate-free ensemble and then asking what distinguishes its members. We combine long unbiased molecular dynamics (MD) with a variational autoencoder (VAE)^23,24^ and VAMPnets^25^ to construct a low-dimensional representation and a Markov state model (MSM)^26–29^ of the apo enzyme, respectively. We then project the dye-bound conformations onto this representation, analyze the correlated motions that connect the two boundary loops, and finally compute the thermodynamic cost of a single point mutation, T114A, within each of the resolved states. This sequence deliberately separates three different kinds of statement: structural overlap, dynamical correlation, and the response to a specific perturbation.

## Results and Discussion

### Machine-learned contact features provide a compact representation of the apo ensemble

Our first task was to find a small set of coordinates that can describe the conformational changes of this 499-residue enzyme. Following the strategy of Ahalawat and coworkers^30^, we trained a random forest (RF) classifier to discriminate between the substrate-free and the five dye-bound ensembles of our earlier work^11^, and used the feature importances to rank the interresidue distances. Out of approximately 1,500 candidate residue pairs, the 38 highest-ranked features already account for 80% of the cumulative impurity-based importance (SI Appendix, Fig. S1 and Table S1), and we have used their Cα–Cα distances for all subsequent clustering, dimensionality reduction and kinetic modeling. Contact-based feature selection of this kind has worked well in other MSM applications^31^. The selected pairs are not confined to the two loops: they are distributed over the binding pocket, the region around the copper centers and several distant parts of the protein (SI Appendix, Fig. S1b), so the classifier is clearly using information well beyond the immediate substrate-contacting region.

Since everything that follows is built on this representation, it is worth stating its scope. The candidate list was generated from spatial contacts in a single apo reference frame, retaining residue pairs separated by more than three positions in sequence whose minimum heavy-atom distance was within 4.5 Å. Such a list reports the rearrangement of existing local packing rather than the formation of entirely new long-range contacts, and it is insensitive to side-chain reorientations that leave the Cα–Cα distances unchanged. The ranking was also supervised by the dye labels, which is worth keeping in mind when we compare bound and apo conformations in these coordinates later.

Density-based clustering (HDBSCAN)^32^ of the six 500-ns ligand-deleted apo segments from our earlier simulations^11^ separates them cleanly into six clusters, and the assignment is unchanged for both tested values of min_samples (SI Appendix, Fig. S3). Separation by trajectory of origin, however, can equally well reflect the persistence of the starting conformation over the accessible simulation time. This is precisely why we generated a much larger set of trajectories, seeded by tICA-metadynamics^33^, before attempting any kinetic analysis.

For the larger dataset (43 μs of unbiased trajectories retained for analysis; see Materials and Methods) we compared two low-dimensional representations: time-lagged independent component analysis (tICA)^34^ and a two-dimensional VAE latent space, both fitted to the same input distances. The six initial apo clusters overlap appreciably in the first two tICA coordinates, whereas the VAE latent coordinates keep them apart (SI Appendix, Fig. S6 a and b), and the implied timescales obtained from the two representations are comparable (SI Appendix, Fig. S6 c and d). We have therefore used the VAE coordinates Z1 and Z2 for visualization throughout, while noting that the cleaner separation is a matter of resolution rather than a claim to a unique reaction coordinate. Fig. 2a shows the free energy landscape projected onto these two coordinates.

**Fig. 2.**
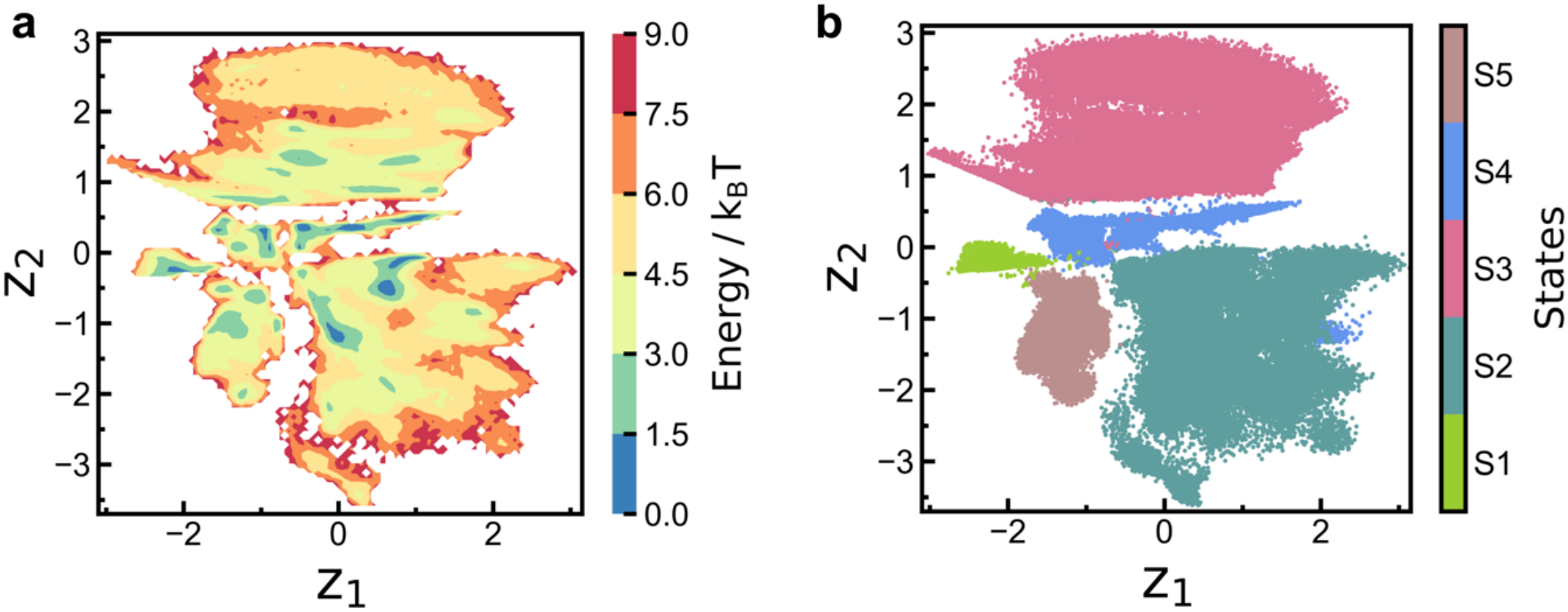
The substrate-free conformational landscape. (a) Free energy surface (in units of k_B_T) of the apo enzyme projected onto the two VAE latent coordinates, Z1 and Z2, obtained from 43 μs of unbiased trajectories. (b) The same projection colored by the five states S1–S5 obtained from the five-output VAMPnets model; the four- and six-state alternatives are shown in SI Appendix, Fig. S8. Relative basin depths reflect the sampled distribution and this projection.

Rather than fixing the number of states in advance, we trained VAMPnets with four, five and six output states on the same data (SI Appendix, Figs. S7 and S8). All three provide acceptable discretizations, and the four- and five-state models carry smaller uncertainties in the slow timescales than the six-state model (SI Appendix, Fig. S8 d–f). We adopt the five-state model, labeled S1–S5, as the working resolution for the structural analysis below (Fig. 2b); the structural picture that emerges is the same at the coarser and finer discretizations, so the specific choice of five states is a matter of convenience.

### The active site loops interconvert between distinct metastable conformations

The MSM built on the five-state discretization gives the kinetic network of Fig. 3. S2 carries the largest stationary population, followed by S4, and the estimated interconversion rates span 1.1–32.5 ms^−1^, the S1 → S4 transition being the fastest. The implied timescales and the Chapman–Kolmogorov test are given in SI Appendix, Figs. S8e and S9^35^. Since these numbers are inferred from a large collection of short, separately initiated trajectories, we read the network as a picture of connectivity and of the approximate separation of timescales rather than as a set of precisely determined rate constants.

**Fig. 3.**
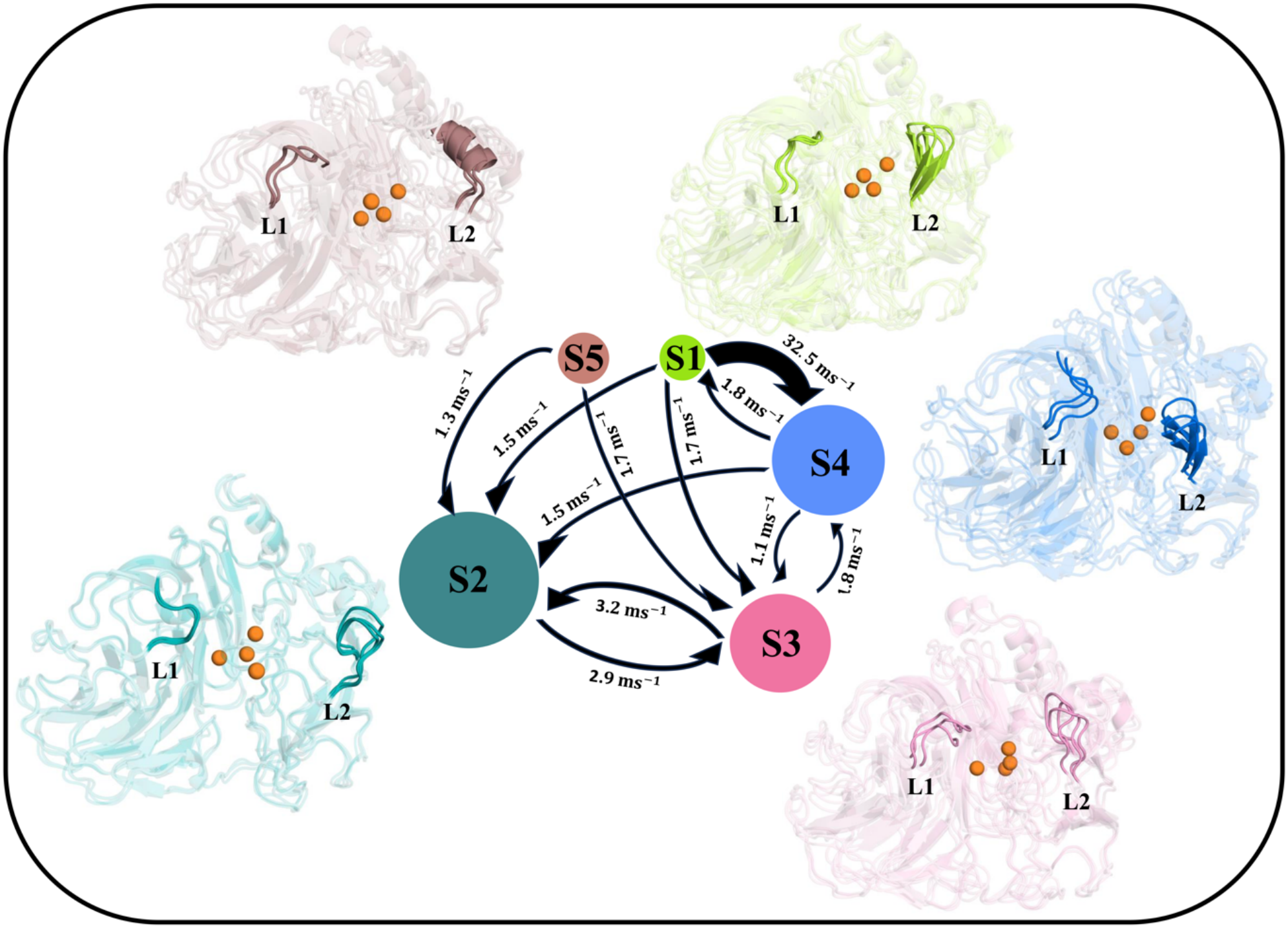
Kinetic network of the five-state model. Circle areas scale with the estimated stationary populations, and the arrows are labeled with the inferred interconversion rates; connections slower than 1 ms^−1^ have been omitted and arrow widths scale with the displayed rates. The representative structures illustrate the conformations of loops L1 and L2 in each state, with copper ions shown as orange spheres. Rates and populations are obtained from the five-state discretization of 43 μs of aggregated trajectories and are meant to convey connectivity and relative timescales.

The structural content of the model is quite striking. The representative structures in Fig. 3 show that the two loops adopt clearly different conformations in the five states. L1 remains largely irregular but changes both its position and its shape, whereas L2 is disordered in S2 and S3, participates in an antiparallel β-sheet in S1 and S4, and folds into a short helix in S5. The volume and the shape of the pocket are therefore different in each state. Thus, the “shapeshifting” behavior that we had earlier observed for L1 alone^11^ turns out to be part of a larger and more cooperative rearrangement involving both loops, and it defines the set of conformational alternatives that an incoming substrate can encounter.

What distinguishes these states at the level of specific interactions? We computed the hydrogen bond occupancies along representative trajectories drawn from each state^36^. A number of residue pairs in and around the binding pocket show markedly different occupancies between the states (SI Appendix, Fig. S10a and Table S2), including several interactions involving L2 and the short helix that carries T114. The N331–F338 hydrogen bond, for example, is present in 96% of the frames in S1 but only 12% in S2, and the T114–D118 side-chain hydrogen bond, to which we return below, varies between 9% and 98% across the five states.

Taken as patterns rather than as individual bonds, the statewise hydrogen bond maps differ by more than 40% in every pairwise comparison (SI Appendix, Fig. S10b), and the contacts that appear or disappear are distributed well beyond the active site (SI Appendix, Fig. S10 c–e). In other words, the loop rearrangement is not a local event: it is accompanied by an extensive rewiring of the intraprotein interaction network, much as we had found for the population shift of the hydrogen-bond network in the PDZ3 domain^37^. Fast hydrogen bond dynamics is known to gate much slower conformational transitions^38^, and the present comparison of state ensembles is a natural starting point for identifying which specific interactions are worth perturbing. That is exactly what we do below for one of them.

### Dye-bound conformations map onto specific regions of the apo landscape

We next projected the five 1-μs dye-bound trajectories of our earlier work^11^ onto the apo VAE coordinates (Fig. 4a). The result is a clean partitioning: brilliant-blue-bound conformations fall in the region of S3; methyl green, coumarin 343 and thioflavin T all map onto S4; and crystal violet maps onto S1. Different dyes are thus accommodated by structurally distinct arrangements of the same pocket, and three of the five apo states – S1, S3 and S4 – have a counterpart among the simulated complexes. We shall refer to these as dye-associated states.

**Fig. 4.**
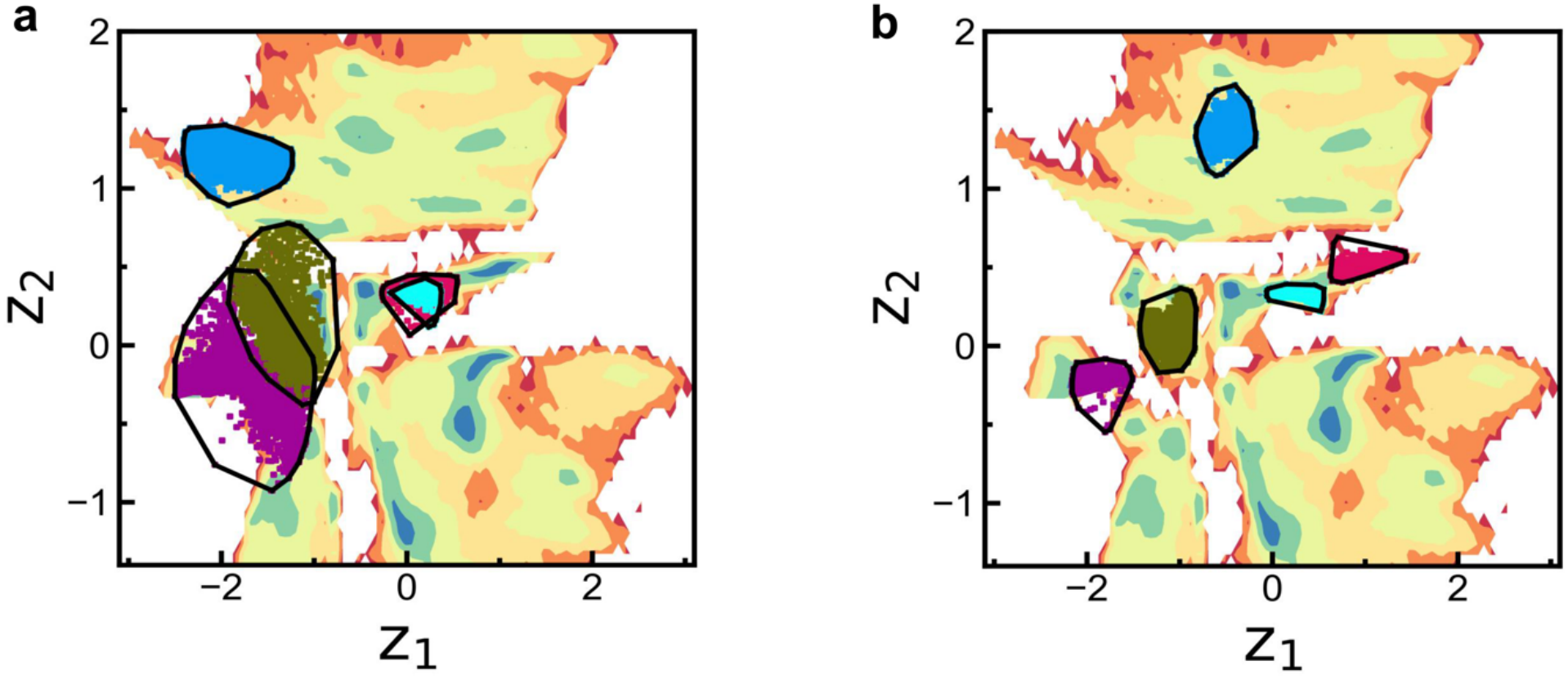
Dye-bound and ligand-deleted trajectories in the apo representation. (a) Projections of the protein conformations sampled in the presence of brilliant blue (blue), coumarin 343 (pink), crystal violet (violet), methyl green (green) and thioflavin T (cyan) onto the apo free energy landscape of Fig. 2a. (b) The last 500 ns of the corresponding trajectories after deletion of the dyes. Black outlines delineate the projected state regions.

When each dye is deleted and the protein is allowed to relax, the last 500 ns of the corresponding trajectory remain largely within the same region of the landscape, the brilliant-blue-deleted trajectory relaxing within S3 (Fig. 4b). The bound-like conformations are therefore not propped up by the ligand: they are metastable in its absence as well, and they belong to the repertoire that the free enzyme visits on its own. This is the observation one expects if binding proceeds through a population shift among pre-existing conformations^39^. Deciding how much of the binding actually follows this route and how much proceeds through induced fit would require simulations of the association process itself, which we have not attempted here.

### A correlation network links the two loops through the T1 copper region

The two loops L1 and L2 sit on opposite sides of the pocket, and their conformations change together. Are their motions correlated, and if so, through which part of the protein? Normalized linear mutual information (LMI)^40,41^ computed over the state-specific segments shows appreciable correlation between the residues of L1 and L2 (Fig. 5a), the strongest coupling being between F332 and N333 of L2 and P163 and L164 of L1. Shortest-path analysis of the correlation graph, with F332 as source and F162 as target, returns a family of closely related routes (SI Appendix, Table S3); the shortest of them is F332–F337–F463–L459–F457–D456–I455–S113–T114–K157–A161–F162 (Fig. 5b). It is worth stressing that this route runs through the protein matrix surrounding the T1 copper and not through the metal ion itself. Among the residues along the path, T114 carries the highest betweenness centrality by a clear margin (Fig. 5c).

**Fig. 5.**
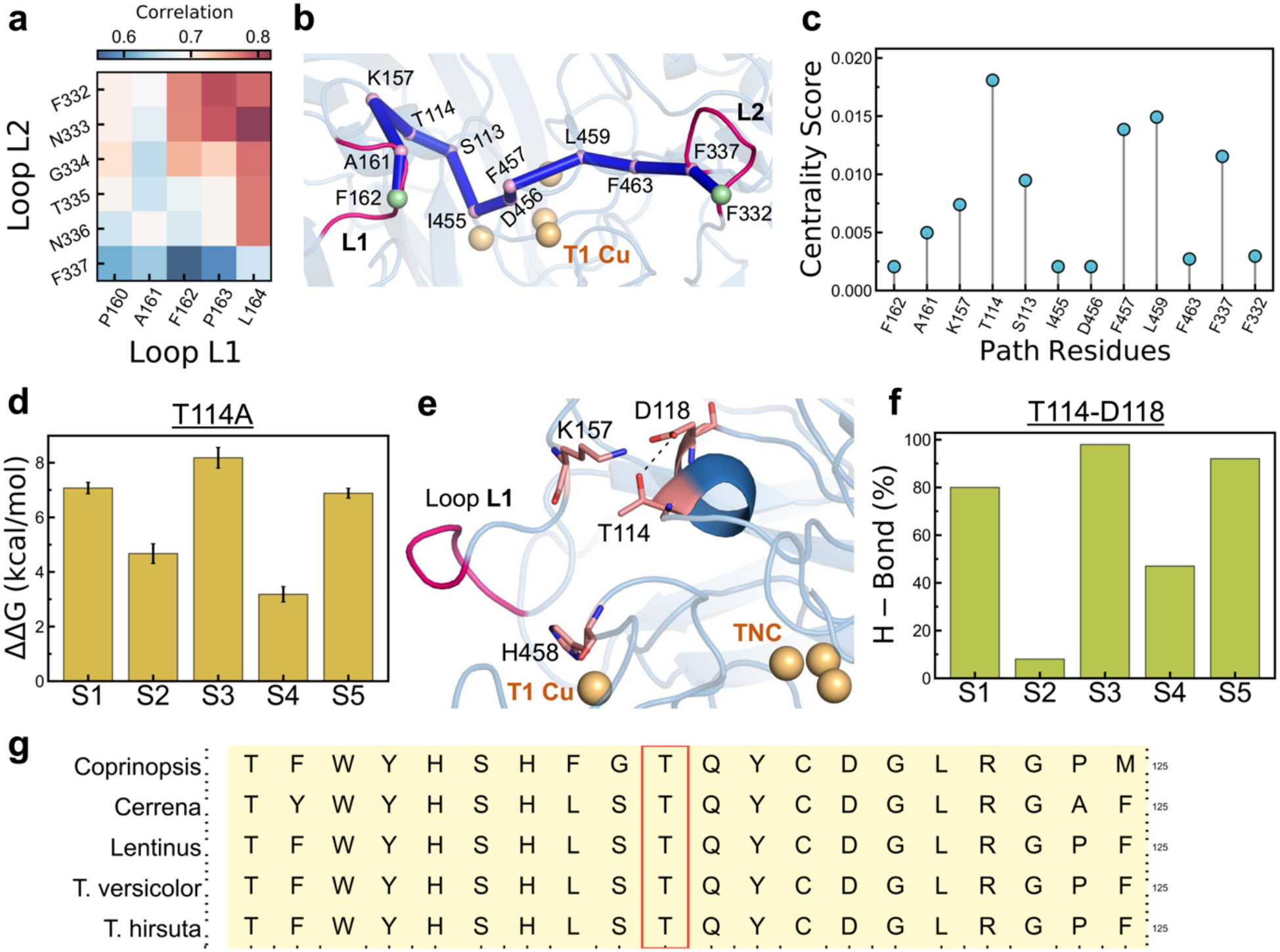
Correlated motions of the two loops and the thermodynamic consequence of T114A. (a) Normalized linear mutual information between the residues of loops L1 and L2. (b) The shortest correlation path connecting F332 (L2) to F162 (L1). Path residues and the two end points are shown as pink and green spheres, respectively, and copper ions in orange. (c) Betweenness centrality of the residues along this path. (d) Mutation-induced destabilization ΔΔG of S1–S5 relative to the common capped-peptide reference, as defined in Eq. 6. Bars are differences of means over five independent replicates per folded state and five peptide replicates; error bars are the propagated standard errors of those means. (e) The T114–D118 side-chain hydrogen bond behind loop L1. (f) Statewise occupancy of the T114–D118 hydrogen bond. (g) Local sequence alignment of five fungal laccase structures, with the threonine corresponding to T114 boxed.

Network measures of this kind are widely used to identify coupled motions and to nominate candidate sites for perturbation^42–46^, and that is how we use them here. They report correlated positional fluctuations and do not by themselves fix the direction in which a perturbation propagates, so we treat T114 as a well-motivated candidate and now ask what happens thermodynamically when its side chain is removed.

### The T114A substitution destabilizes the five states to very different extents

T114 lies near the N-terminal cap of a short helix adjacent to L1 and donates a side-chain hydrogen bond to D118 (Fig. 5e). To test its role we removed the hydroxyl-bearing side chain *in silico* by nonequilibrium alchemical mutation to alanine^47,48^, computing the mutation free energy separately in each of the five folded states and in a capped ACE–GTG–NME reference peptide (Materials and Methods and Fig. 6). A positive value of the resulting ΔΔG means that the mutation costs more in the folded state than in the reference, that is, it is destabilizing.

**Fig. 6.**
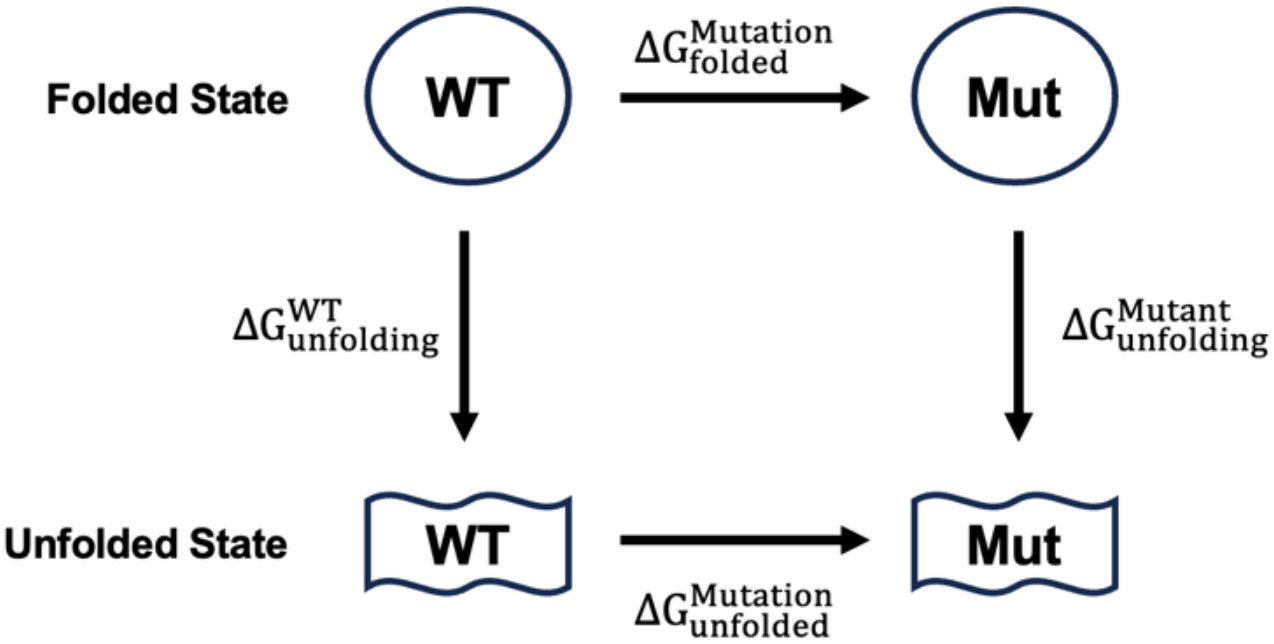
Thermodynamic cycle used for T114A. The horizontal legs connect the wild type and the mutant, and the vertical legs connect the folded protein with the unfolded reference, which in our calculations is represented by the capped peptide ACE–GTG–NME and its alanine analogue. The difference between the folded-state and peptide mutation free energies defines ΔΔG in Eq. 6.

Averaged over five fully independent replicates, ΔΔG is 7.06 ± 0.34, 4.67 ± 0.21, 8.17 ± 0.25, 3.18 ± 0.25 and 6.89 ± 0.24 kcal mol^−1^ for S1–S5, respectively (Fig. 5d; SI Appendix, Tables S4 and S5). The spread across the states is the significant result here. Since the common peptide contribution cancels exactly when two states are compared, the differences between states are independent of that reference: S3 is destabilized by 4.99 ± 0.29 kcal mol^−1^ more than S4, and S1 by 3.88 ± 0.37 kcal mol^−1^ more than S4. A difference of 5 kcal mol^−1^ at 300 K corresponds to a shift of more than three orders of magnitude in the relative populations of the two states. Although the absolute values are subject to force-field and sampling error, the calculation predicts that a single side chain can strongly redistribute the conformational ensemble of the active site.

The ranking of the penalties follows the T114–D118 hydrogen bond occupancy rather closely. The three states in which the bond is essentially persistent – S1, S3 and S5, with occupancies of 80%, 98% and 92% – pay the larger cost, while S2 and S4, with occupancies of 9% and 47%, pay much less (Fig. 5f). The correspondence is not strictly monotonic: S4 has a higher occupancy than S2 but a smaller penalty, and S5 a higher occupancy than S1 with a marginally smaller one. This is only to be expected, since a Thr → Ala substitution removes more than one hydrogen bond: it also alters the local packing, the helix capping and the solvation of that region, and helix capping interactions are themselves known to contribute substantially to stability^49^.

The pattern of destabilization is also instructive in what it does not track. S4, which accommodates three of the five dyes, is the least affected state, whereas S5, which is associated with none of them, is strongly destabilized. T114A is therefore not a mutation that simply removes the dye-binding conformations; what it does is tilt the apo ensemble away from S1, S3 and S5 and towards S2 and S4. This redistribution is the prediction that experiments should look for, and translating it into a change of substrate preference will additionally require the effect of the mutation on the bound states and on the chemical steps of turnover. We propose T114A as the primary test, with D118A and a control substitution elsewhere on the same helix to establish the specificity of the interaction.

Two further observations support the choice of this residue. First, T114–F457 and T114–A156 appear independently among the 38 RF-selected features (SI Appendix, Fig. S12 and Table S1); both report the position of the T114 helix relative to the pocket, so the supervised classifier and the correlation analysis arrive at the same region by entirely different routes. Second, the threonine is conserved at the aligned position in all five fungal laccase structures that we examined: *T. versicolor* (1GYC), *Lentinus tigrinus* (2QT6), *Cerrena* sp. (5Z1X), *T. hirsuta* (3FPX) and *Coprinopsis cinerea* (formerly *Coprinus cinereus*; 1A65)^50–54^ (Fig. 5g).

### Scope of the present analysis

A few features of the analysis define the scope of these conclusions. The representation was built from contacts in a single reference frame and ranked with the help of the dye labels, so the correspondence between bound and apo conformations is best read as structural rather than as an independent test of the binding mechanism; the 43-μs dataset is an aggregate of many short trajectories seeded from metadynamics, which constrains the precision of the estimated rates and stationary populations; and fixed protonation states together with a bonded model for the copper centers place the chemical steps of catalysis outside the reach of these simulations. Within this scope, what we report is a physically interpretable picture of the conformational ensemble of one fungal laccase, together with a specific and falsifiable prediction that follows from it.

## Conclusions

The substrate-free simulations of *T. versicolor* laccase resolve several distinct arrangements of the two loops that form the rim of the binding pocket, and show that the rearrangement of these loops is accompanied by an extensive rewiring of the hydrogen-bond network beyond the active site. Conformations sampled with five different bound dyes occupy specific regions of this landscape, so each dye is accommodated by a conformation that the free enzyme already visits. A correlation network connects the two loops through the protein region surrounding the T1 copper and identifies T114 as the residue with the highest betweenness along that route. Nonequilibrium alchemical mutation calculations then show that removing this single side chain destabilizes the five conformers by amounts differing by up to 5 kcal mol^−1^, large enough to strongly shift their relative populations under the simulation model. The conformational balance of a promiscuous active site can therefore be tuned through a distant residue that never directly interacts with substrates, and the obvious next step is a direct comparison of the wild type and the T114A mutant, in which the redistribution predicted here can be confronted with measured binding and turnover.

## Materials and Methods

### Molecular dynamics simulations

All simulations start from the crystal structure of *T. versicolor* laccase bound to 2,5-xylidine (PDB 1KYA)^14^. The protein was described by the ff14SB force field^55^ and the copper centers by the bonded model generated with MCPB.py^56^ in AmberTools20^57^, exactly as in our earlier work^11^. All four copper ions were retained in the fully oxidized Cu(II) state in every simulation. The earlier dataset consists of five 1-μs dye-bound trajectories and six 1-μs ligand-deleted trajectories, one for each of the five dyes and one for 2,5-xylidine; the second half of each ligand-deleted trajectory (500 ns) enters the apo analysis.

To this we added 100 unbiased 500-ns apo trajectories started from structures selected along the tICA-metadynamics^33^ runs, giving 50 μs of new production. Discarding the first 100 ns of each as equilibration, we retained 400 ns from each of them (40 μs) and combined these with the six earlier 500-ns segments (3 μs) to obtain the 43-μs dataset used for all apo analyses. The 5 μs of dye-bound trajectories were used only for feature selection and for the structural projections of Fig. 4, and the metadynamics trajectories only to generate starting structures; neither contributes to the kinetic estimates.

Each solvated system contained approximately 17,000–20,000 TIP3P^58^ water molecules and 14 Na^+^ counterions. All simulations were performed with GROMACS 2024.2^59^. After steepest-descent energy minimization and NVT and NPT equilibration, production runs were carried out at 300 K and 1 atm using the velocity-rescaling thermostat^60^ and the Parrinello–Rahman barostat^61^. Periodic boundary conditions were applied in all directions, long-range electrostatics was treated by particle mesh Ewald^62^, and the real-space cutoff for electrostatic and van der Waals interactions was 1.0 nm. Bonds involving hydrogen atoms were constrained with LINCS^63^, the integration time step was 2 fs, and coordinates were saved every 100 ps.

### Random forest feature selection

The RF dataset consisted of the five 1-μs dye-bound trajectories and the last 500 ns of the xylidine-deleted apo trajectory, with each frame labeled by its trajectory of origin (six classes). Candidate residue pairs were defined in a single representative frame of the apo trajectory: pairs separated by more than three positions in sequence were retained if their minimum heavy-atom distance was 4.5 Å or less, which gave approximately 1,500 pairs. Their Cα–Cα distances, computed with MDTraj^64^, formed the input to the classifier. Note that the 4.5 Å criterion defines the candidate list only; it is not a cutoff imposed on the Cα–Cα distances themselves, which vary freely during the trajectories.

A scikit-learn RF classifier^65,66^ with 10 trees was trained on a randomly selected 70% of the frames and tested on the remaining 30%. The Gini impurity measures the class mixing within a node, and the importance of a feature is the decrease in impurity attributed to it, weighted by the number of samples reaching each split, averaged over the trees and normalized over all features. Ranking the features by this quantity and retaining those that account for 80% of the cumulative importance left 38 residue pairs. The validation accuracy was 0.998; since neighboring frames are correlated and the class labels coincide with individual trajectories, this figure measures discrimination within the sampled dataset. The stability of the selected set under 200 repetitions of the random split is shown in SI Appendix, Fig. S2.

### Density-based clustering and enhanced sampling

HDBSCAN^32^ was applied to the selected-distance representation of the six 500-ns apo segments. We compared min_samples values of 1,000 and 2,000 (shown in SI Appendix, Fig. S3) to investigate how changing the local cluster density affects the final results.

tICA^34^ was fitted to the same six segments with a 20-ns lag using PyEMMA 2.5.7^67^, and the first four components were used as collective variables for tICA-metadynamics^33^, performed with GROMACS 2021.5 patched with PLUMED 2.7.4^68^. Each metadynamics run was 100 ns long, with Gaussian hills of initial height 1.2 kJ mol^−1^ and width 0.5 (in units of the biased tICA coordinates) deposited every 500 steps and a bias factor of 8. One hundred structures spanning the sampled region were then selected as starting points for the unbiased runs (SI Appendix, Fig. S4 d–f). A separate tICA fit, with the same 20-ns lag, was also performed on the complete 43-μs apo dataset for comparison (SI Appendix, Fig. S4 g–i).

### Variational autoencoder

The VAE^23^ was implemented in PyTorch and trained on the 38 selected distances from the 43-μs apo dataset. The encoder consists of fully connected hidden layers with 300, 200, 100 and 20 nodes followed by a two-dimensional latent representation, and the decoder mirrors this architecture; the first hidden layer uses tanh and the remaining ones ReLU activation. The loss function (Eq. 1) combines a mean-squared reconstruction term (Eq. 2) with the Kullback–Leibler regularization^69^ (Eq. 3):

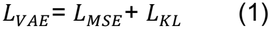

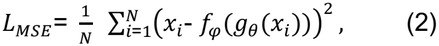

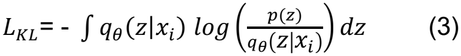

Here x_i_ is an input vector, z the latent variable, g_θ_ the encoder and f_φ_ the decoder; the reconstruction term is evaluated with samples drawn from the encoder distribution q_θ_(z|x_i_), and p(z) is the standard normal prior. For a Gaussian encoder, the regularization term for each input reduces to

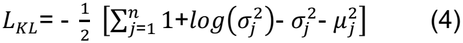

where n is the latent dimension and μ_j_ and σ_j_^2^ are the corresponding means and variances. The distances were standardized using the scikit-learn StandardScaler. We used a 70:30 training–validation split, 20 epochs, a batch size of 500, and the Adam optimizer^70^ with a learning rate of 10^−5^; the training and validation losses are shown in SI Appendix, Fig. S5.

### VAMPnets and Markov state modeling

VAMPnets^25^ were implemented using Deeptime^71^ on the same 43-μs dataset. Standardized instantaneous and time-lagged feature vectors were paired at a training lag of 350 ns. Each network lobe has three fully connected hidden layers with 36, 24 and 16 nodes and ReLU activation, and a softmax output layer with four, five or six nodes providing the alternative state assignments. The models were trained for 100 epochs on 80% of the data with 20% held out for validation, a batch size of 500 and the Adam optimizer with a learning rate of 10^−4^. The VAMP-2^72^ training and validation scores are shown in SI Appendix, Fig. S7.

For an MSM, K_ij_(τ) is the probability of finding the system in state j after a lag time τ, given that it started in state i, and the implied timescale associated with a positive non-stationary eigenvalue λ_i_ of the transition matrix is t_i_ = −τ/ln λ_i_. Approximate independence of t_i_ from the lag time is the usual consistency check^35^. We estimated the transition matrix on the five-state discretization at a lag of 70 ns using Deeptime. The 350-ns VAMPnets training lag and the 70-ns transition-matrix lag serve different purposes and are not interchangeable. The lag dependence of the timescales and the Chapman–Kolmogorov comparison are given in SI Appendix, Figs. S8 and S9.

### Hydrogen bond and residue correlation analyses

Hydrogen bond occupancies were computed along representative trajectories assigned to each apo basin, counting both backbone and side-chain interactions with the usual geometric criteria (donor–acceptor distance below 3.0 Å and deviation of the donor–H–acceptor angle from linearity below 20°). For a residue pair capable of forming more than one hydrogen bond the summed occupancy can exceed 100% (SI Appendix, Table S2). For the statewise pattern comparison of SI Appendix, Fig. S10 we retained pairs with occupancy above one-third in at least one state.

Residue-position fluctuations were analyzed with the correlationplus package^41^ on five 5-ns segments, one drawn from each apo state. The normalized Gaussian linear mutual information between the positions of residues i and j is^40^

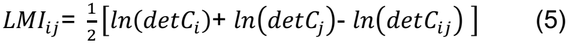

where 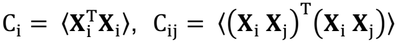 and **X**_i_ = **R**_i_ − ⟨**R**_i_⟩, where **R**_i_ is the position vector of the i^th^ Cα atom. Shortest paths were computed on the resulting correlation graph with F332 as source and F162 as target.

### Nonequilibrium alchemical mutation

We define the mutation-induced destabilization as the difference between the unfolding free energies of the wild type and the mutant, which the thermodynamic cycle in Fig. 6 converts into a difference of two mutation free energies:

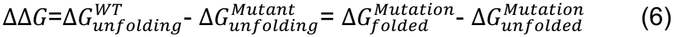

A positive ΔΔG therefore means that the mutation destabilizes the protein. We approximated the unfolded reference by the capped tripeptide ACE–GTG–NME, mutating its central threonine to alanine, and computed the folded-state mutation free energies separately for states S1–S5.

Hybrid structures and topologies were generated with pmx^73^ using its amberff14sb-mutant force field, supplemented with the same MCPB-derived copper parameters as the baseline simulations, and the alchemical simulations were run with GROMACS 2022.5. The wild-type and mutant end states were equilibrated separately for 20 ns in TIP3P water at 300 K and 1 atm following energy minimization, and the folded end states were checked to remain within their assigned basins. One hundred starting configurations were extracted uniformly from the final 18 ns, and each was used to launch a 100-ps switching trajectory to the opposite end state. The entire procedure, from equilibration to switching, was repeated five times independently for every folded state and for the peptide.

The alchemical Hamiltonian H(x;λ) connects the two end states, with λ = 0 for the wild type and λ = 1 for the mutant; schematically,

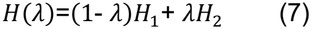

The actual interpolation of the nonbonded interactions, including the soft-core treatment, is defined by the simulation input. During a 100-ps switch with a 2-fs time step, λ changes by +2 × 10^−5^ per step in the forward direction and by −2 × 10^−5^ per step in reverse. The work accumulated along a switching trajectory is

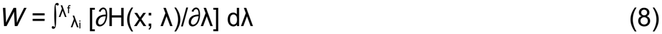

where λ_i_ and λ_f_ are 0 and 1 for the forward switches and 1 and 0 for the reverse ones. W is the total protocol work, the dissipated work being W minus the equilibrium free energy difference for that direction. The forward and reverse work distributions are related by the Crooks fluctuation theorem^48^,

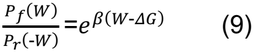

with β = 1/k_B_T, where P_f_(W) is the forward and P_r_(−W) the sign-reversed reverse work distribution. We obtained the equilibrium free energy difference from both work samples using the Bennett acceptance ratio (BAR) estimator^74^, which provides the maximum-likelihood solution implied by Eq. 9.

The work distributions for all systems and replicates are shown in SI Appendix, Fig. S11, and the individual free energy estimates with their within-replicate BAR uncertainties are listed in SI Appendix, Table S4. For the summary values we weighted the five independent replicates equally and took the standard error. For ΔΔG we propagated the standard errors of the folded-state and peptide means in quadrature (SI Appendix, Table S5). Since all five ΔΔG values share the same peptide mean, this reference uncertainty is common to them and cancels when two folded states are compared.

## Acknowledgments

We thank the Anusandhan National Research Foundation (ANRF), Government of India, for support through projects MTR/2023/000336 and ANRF/ARGM/2025/002124/TS, which enabled the analyses reported here. We also acknowledge the PARAM Rudra supercomputing facility at the S. N. Bose National Centre for Basic Sciences, established under the National Supercomputing Mission, Government of India, for computational resources. S.M. thanks the S. N. Bose National Centre for Basic Sciences for a research fellowship. OpenAI Codex and Claude Opus large language models were used solely for language editing and consistency checks during revision. The authors reviewed and verified all text and remain responsible for the scientific content.

## Author contributions

S.C. and R.B. designed research; S.M. performed research; S.M., R.B. and S.C. analyzed data; and S.M. and S.C. wrote the paper.

The authors declare no competing interest.

## Data availability

The representative structures of the five states, the selected-distance trajectories, the trained VAE and VAMPnets models, the MSM count and transition matrices, the copper-center parameters, the alchemical inputs and work data, and all analysis scripts will be deposited in Zenodo before publication, and the persistent identifier and dataset citation will be added here. These materials are available from the corresponding authors for editorial and peer review. All other data are included in the manuscript and SI Appendix.

