## Supplementary Information for "Conformational heterogeneity and state-dependent stability of active-site loops in *Trametes versicolor* laccase"

### SI Appendix

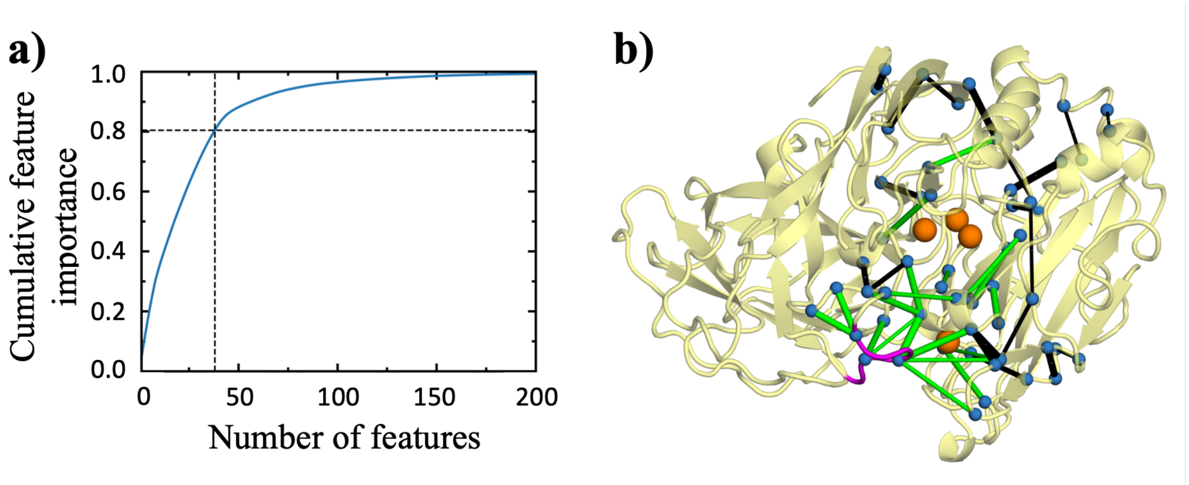


**Fig. S1.** Random forest feature selection. **(a)** Cumulative feature importance for the 200 highest-ranked residue pairs; the dashed lines mark the 38 features that together account for 80% of the total importance. **(b)** The 38 selected pairs drawn as connections between Cα atoms (blue spheres). Green connections involve residues of loop L1 (G159–L164) and the catalytic region, and black connections denote the remaining selected pairs; line widths scale with the RF feature importance. Loop L1 is shown in magenta and the copper ions in orange.

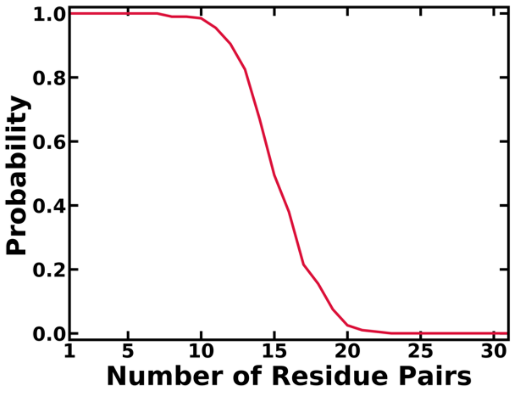


**Fig. S2.** Stability of the random forest feature selection under 200 random splits of the data. The probability is that of obtaining n different residue pairs when the 38 pairs selected in the primary analysis are compared with the pairs satisfying the same criteria in replicate random forest runs. Since neighboring frames are correlated and the class labels coincide with individual trajectories, this test probes the stability of the ranking within the sampled dataset.

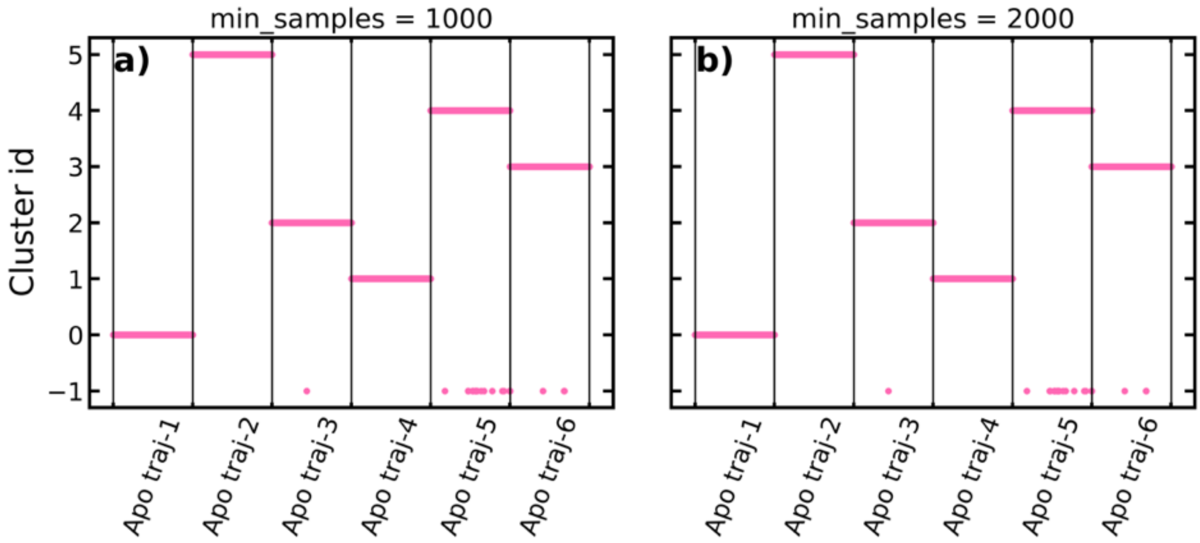


**Fig. S3.** HDBSCAN cluster labels for the last 500 ns of each of the six ligand-deleted apo trajectories, obtained with min_samples = 1,000 **(a)** and 2,000 **(b)**. Each group along the abscissa is one trajectory and the ordinate gives the cluster label; points labeled −1 are unassigned. This parameter controls the local cluster density.

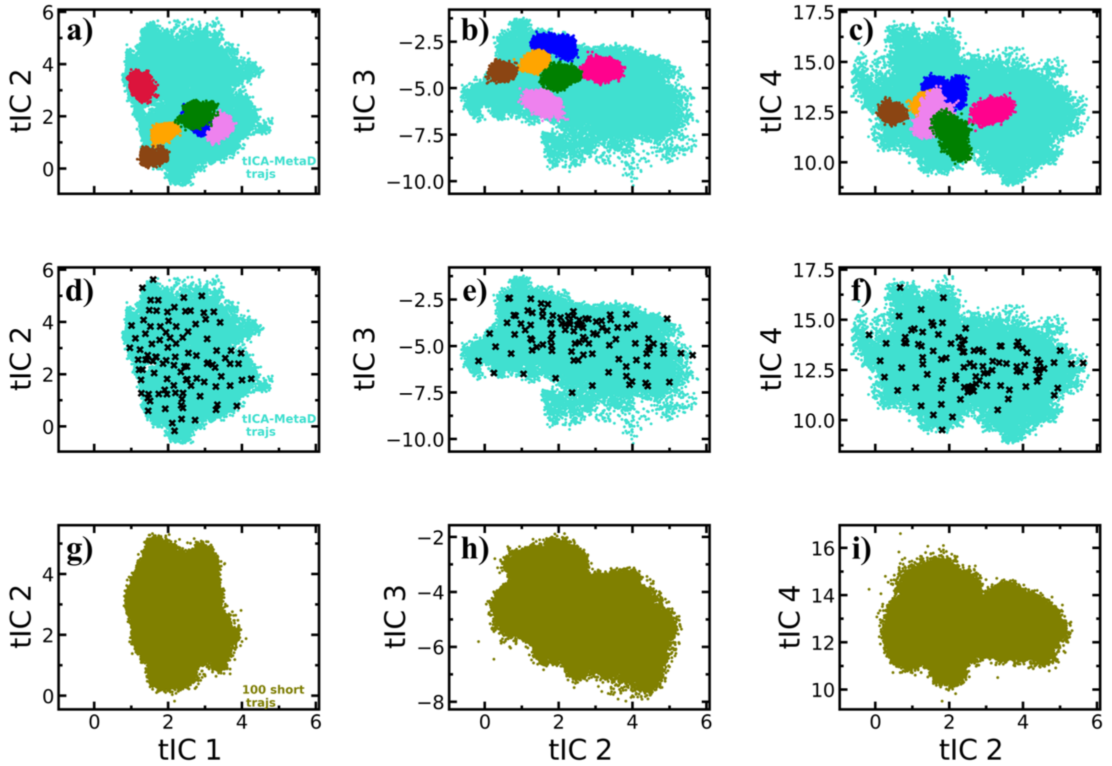


**Fig. S4.** tICA projections and selection of the starting structures. **(a–c)** The last 500 ns of the six ligand-deleted trajectories, colored as 2,5-xylidine (red), brilliant blue (blue), coumarin 343 (orange), thioflavin T (brown), methyl green (green) and crystal violet (violet), shown together with the tICA-metadynamics frames (cyan). **(d–f)** Crosses mark the 100 structures selected to seed the unbiased trajectories. **(g–i)** The 100 unbiased apo trajectories projected on the new tICA coordinates; each run was 500 ns long and 400 ns of each was retained for analysis.

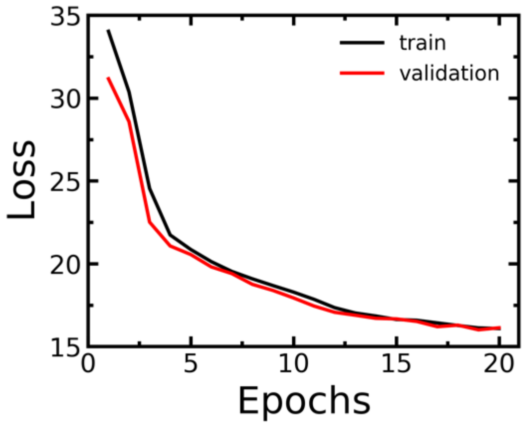


**Fig. S5.** Training and validation losses of the variational autoencoder over 20 epochs.

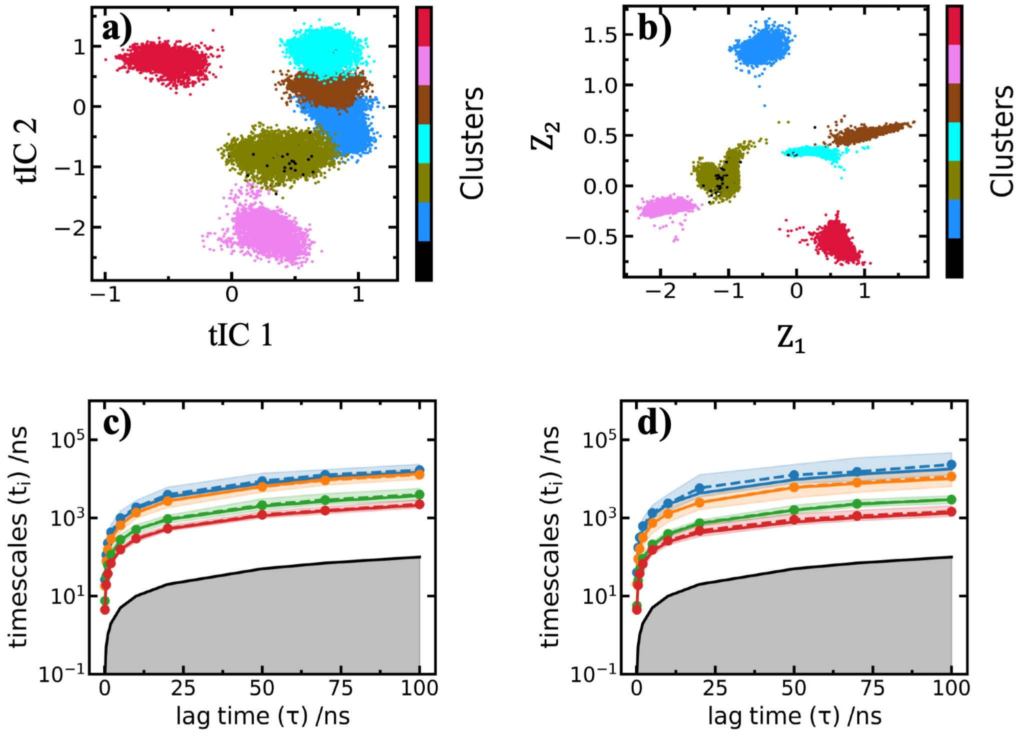


**Fig. S6.** Comparison of the tICA and VAE representations. **(a, b)** The six ligand-deleted apo segments shown in the first two tICA coordinates and in the two VAE latent coordinates, colored by the HDBSCAN cluster labels. **(c, d)** Implied timescales obtained after k-means discretization in the tICA and in the VAE representation; the bands denote the estimated standard errors and the gray region marks timescales at or below the lag time.

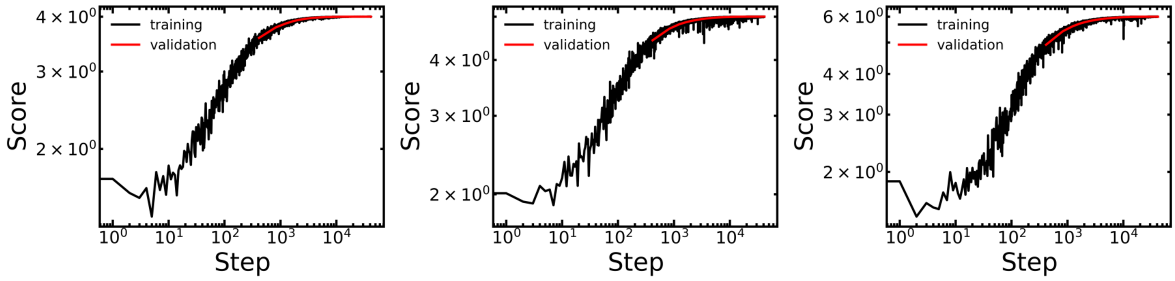


**Fig. S7.** VAMP2 scores of the VAMPnets with four **(a)**, five **(b)** and six **(c)** output states as a function of the optimization step. Black and red curves are the training and the validation scores.

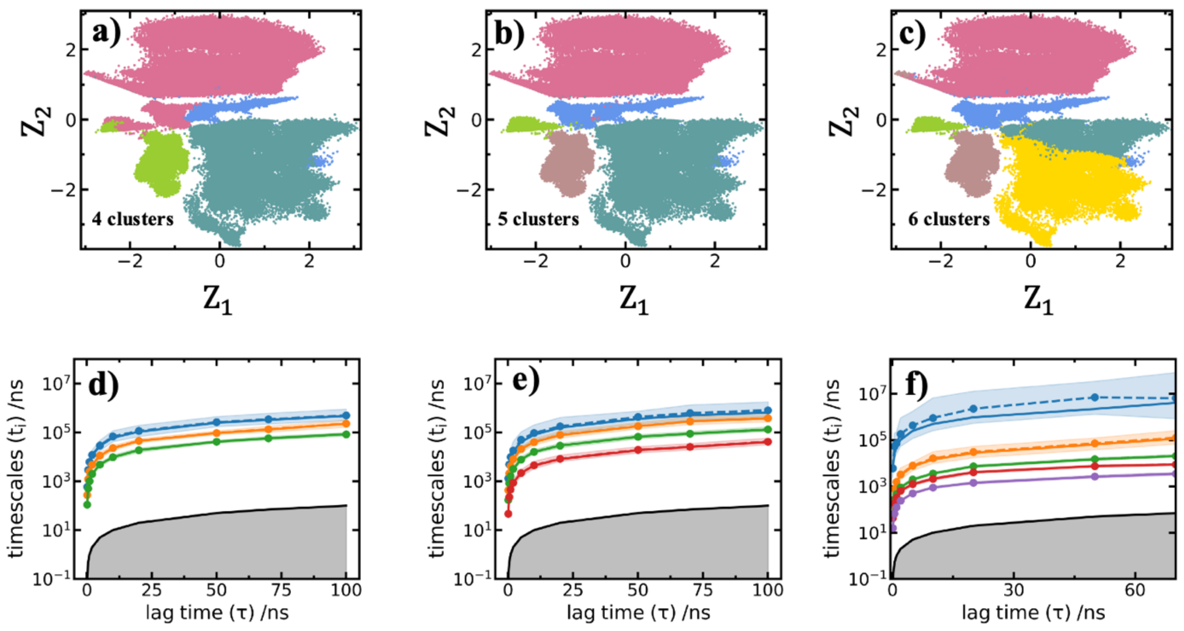


**Fig. S8.** Alternative VAMPnets discretizations and the corresponding implied timescales. **(a–c)** Four-, five- and six-state assignments shown on the VAE latent coordinates. Each color represents a different state. **(d–f)** Implied timescales for these three discretizations; the bands denote the estimated standard errors and the gray regions mark timescales at or below the lag time.

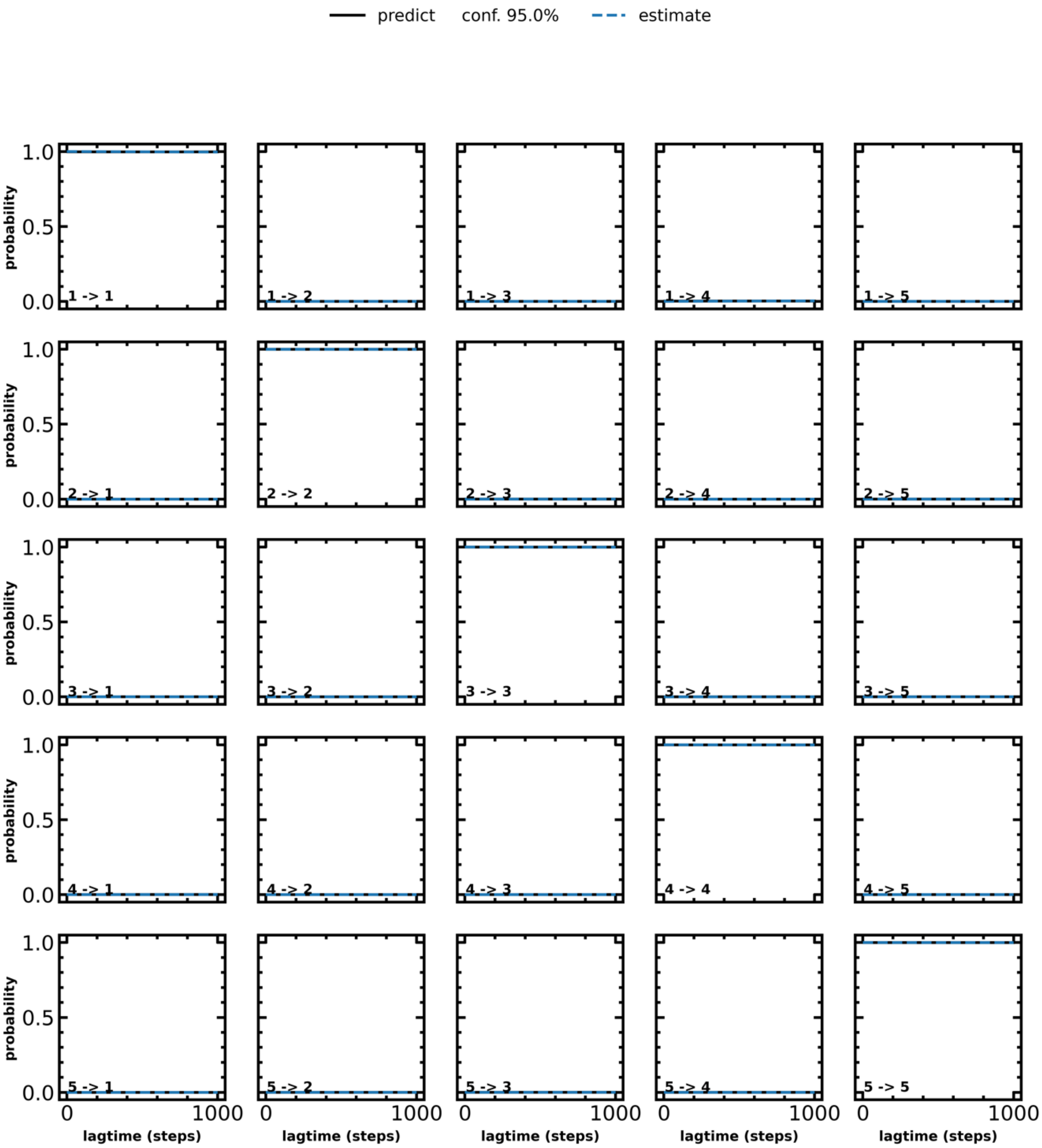


**Fig. S9.** The Chapman–Kolmogorov test for the five-state MSM. Each panel compares the directly estimated transition probability (estimate) with the prediction obtained by propagating the transition matrix (model); the shaded bands are the 95% confidence intervals.

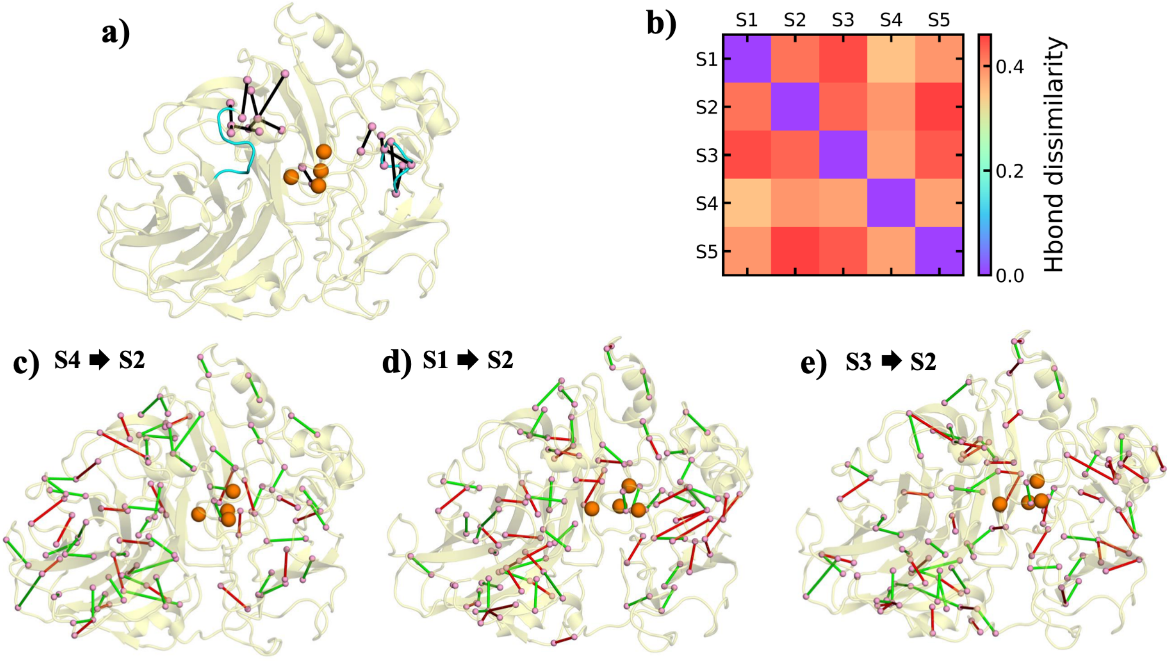


**Fig. S10.** Statewise hydrogen bonding patterns. **(a)** Residue pairs near the substrate binding pocket whose occupancies differ markedly among the five metastable states, S1–S5 (Table S2); loops L1 and L2 are shown in cyan. **(b)** Pairwise dissimilarity of the statewise occupancy patterns. **(c–e)** For the indicated pairs of states, contacts present in the first state and absent in the second are drawn in red, and those present in the second and absent in the first in green.

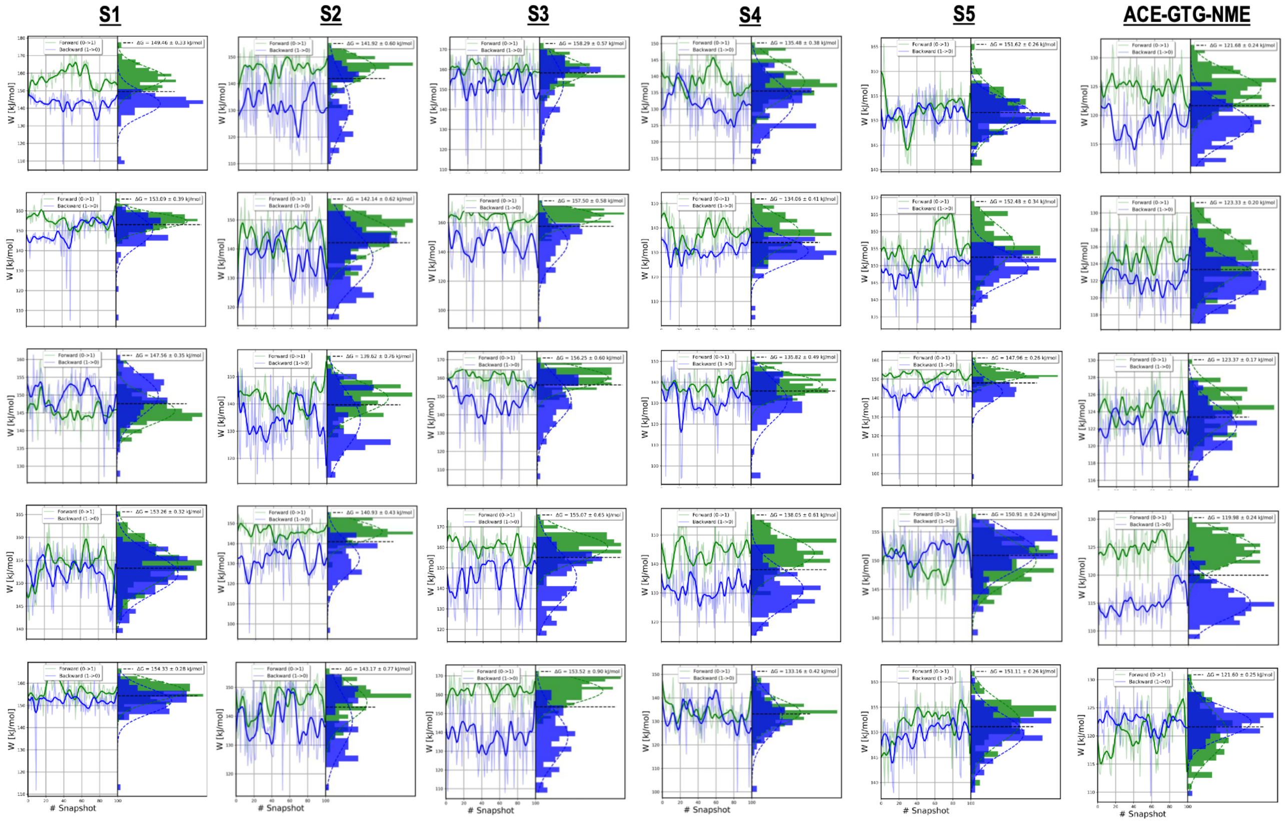


**Fig. S11.** Forward (green) and reverse (blue) nonequilibrium work samples for the T114A mutation in states S1–S5 and in the ACE–GTG–NME reference peptide. Rows correspond to five independent replicates and each distribution contains 100 switching trajectories; the work is given in kJ mol⁻¹. Free energy differences were estimated with the BAR estimator and are listed in Tables S4 and S5.

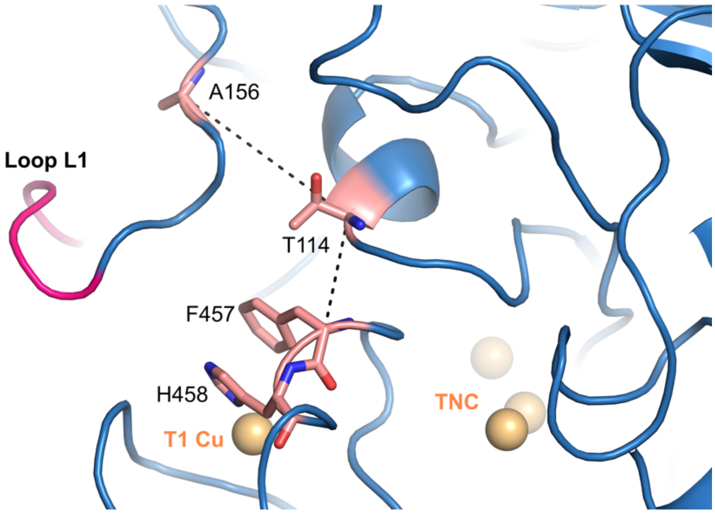


**Fig. S12.** Structural context of the two random-forest-selected features that involve T114. The dashed black lines mark the T114–F457 and T114–A156 Cα–Cα distances. Loop L1 is shown in magenta and the copper ions in orange.

**Table S1.** The 38 residue pairs retained by the random forest feature selection. Their Cα–Cα distances were used for all clustering, dimensionality reduction and kinetic modeling reported in the main text.

| | | | |
|---|---|---|---|
| F337 – A461 | T105 – S225 | K157 – E460 | Q70 – W449 |
| F44 – V126 | P163 – D206 | V425 – H454 | T114 – F457 |
| F162 – F457 | H66 – R243 | F162 – F337 | V154 – G159 |
| P4 – G38 | A390 – H395 | F337 – F344 | A156 – F457 |
| G401 – N445 | H402 – W449 | A155 – G159 | I356 – A475 |
| F397 – F404 | C117 – A156 | P163 – F457 | T114 – A156 |
| Q70 – F97 | W65 – Q70 | F337 – H395 | F44 – V99 |
| F162 – A461 | N336 – A461 | P446 – N478 | F162 – G334 |
| H452 – F463 | F330 – T428 | I82 – F344 | |
| A329 – I339 | F450 – L459 | M328 – N340 | |

**Table S2.** Hydrogen bond occupancies (%) for selected residue pairs around the substrate binding pocket in the five metastable states. A value above 100% indicates that more than one hydrogen bond can form simultaneously between that residue pair, the entry then being 100 times the mean number of bonds. Dashes represent values below the threshold of one-third.

| H-bonding pairs | State S1 | State S2 | State S3 | State S4 | State S5 |
|---|---|---|---|---|---|
| N331-F338 | 96.0 | - | 56.0 | 61.0 | - |
| N331-T335 | - | - | - | - | 70.0 |
| N333-N336 | 43.0 | 84.0 | 131.0 | 51.0 | - |
| F337-S343 | - | - | 71.0 | - | - |
| F344-A464 | - | 53.0 | - | - | - |
| C453-D456 | - | 88.0 | - | - | - |
| T114-D118 | 80.0 | - | 98.0 | 47.0 | 92.0 |
| Q115-L120 | 43.0 | - | 41.0 | - | 43.0 |
| M57-Q115 | - | 45.0 | 57.0 | - | - |
| S110-Q115 | 53.0 | 47.0 | - | - | - |
| S60-Q115 | 63.0 | 59.0 | 49.0 | 33.0 | 57.0 |
| V25-D118 | 35.0 | 49.0 | - | - | - |
| D118-A156 | - | 60.0 | - | - | - |

| | | | | | |
|---|---|---|---|---|---|
| T56-K157 | 39.0 | - | - | - | - |

**Table S3.** Correlation paths connecting F332 in loop L2 (source) to F162 in loop L1 (target), arranged in order of increasing length. The five shortest paths share almost the same set of residues, and T114 appears in every one of them.

| Pathways | Residues involved |
|---|---|
| Path-1 | F332 – F337 – F463 – L459 – F457 – D456 – I455 – S113 – **T114** – K157 – A161 – F162 |
| Path-2 | F332 – F337 – F463 – L459 – F457 – D456 – I455 – S113 – **T114** – A156 – K157 – A161 – F162 |
| Path-3 | F332 – F337 – F463 – L459 – F457 – D456 – I455 – S113 – **T114** – K157 – L158 – P160 – A161 – F162 |
| Path-4 | F332 – F337 – F463 – L459 – F457 – D456 – I455 – S113 – **T114** – A156 – K157 – L158 – P160 – A161 – F162 |
| Path-5 | F332 – N336 – F337 – F463 – L459 – F457 – D456 – I455 – S113 – **T114** – K157 – A161 – F162 |

**Table S4.** Mutation free energies (kcal mol⁻¹) from five independent replicates, for the folded states S1–S5 and for the capped peptide reference. Each replicate value is given with its within-replicate BAR uncertainty, and the final row gives the unweighted mean ± standard error of the mean obtained from the scatter of the five independent replicate estimates. The capped peptide serves as a common reference for all five states.

| State S1 | State S2 | State S3 | State S4 | State S5 | ACE-GTG-NME |
|---|---|---|---|---|---|
| $\Delta G_{\text{folded}}^{\text{Mutation}}$ | | | | | $\Delta G_{\text{unfolded}}^{\text{Mutation}}$ |
| $35.72 \pm 0.08$ | $33.92 \pm 0.14$ | $37.83 \pm 0.14$ | $32.38 \pm 0.09$ | $36.24 \pm 0.06$ | $29.08 \pm 0.06$ |
| $36.59 \pm 0.09$ | $33.97 \pm 0.15$ | $37.64 \pm 0.14$ | $32.04 \pm 0.10$ | $36.43 \pm 0.08$ | $29.48 \pm 0.05$ |
| $35.27 \pm 0.08$ | $33.37 \pm 0.18$ | $37.43 \pm 0.14$ | $32.46 \pm 0.12$ | $35.36 \pm 0.06$ | $29.49 \pm 0.04$ |
| $36.63 \pm 0.08$ | $33.68 \pm 0.10$ | $37.06 \pm 0.16$ | $32.99 \pm 0.15$ | $36.07 \pm 0.06$ | $28.68 \pm 0.06$ |
| $36.89 \pm 0.07$ | $34.22 \pm 0.18$ | $36.69 \pm 0.22$ | $31.83 \pm 0.10$ | $36.12 \pm 0.06$ | $29.06 \pm 0.06$ |
| **36.22 ± 0.31** | **33.83 ± 0.14** | **37.33 ± 0.20** | **32.34 ± 0.20** | **36.04 ± 0.18** | **29.16 ± 0.15** |

**Table S5.** State-specific destabilization ΔΔG (kcal $mol^{-1}$), defined in Eq. 6 as the folded-state mutation free energy minus that of the capped peptide and calculated as differences of the means in Table S4. The uncertainties are the standard errors of the means propagated in quadrature. Since the peptide reference is common to all five states, its uncertainty cancels in pairwise comparisons between folded states.

| State S1 | State S2 | State S3 | State S4 | State S5 |
|---|---|---|---|---|
| 7.06 ± 0.34 | 4.67 ± 0.21 | 8.17 ± 0.25 | 3.18 ± 0.25 | 6.89 ± 0.24 |
